# Antibiotic use in early life subsequently impairs MAIT cell-mediated immunity

**DOI:** 10.1101/2024.05.10.593643

**Authors:** Adam L. Sobel, Jonathan Melamed, Dominic Haas, Gabrielle LeBlanc, Aiko Cirone, Michael G. Constantinides

## Abstract

Mucosal-associated invariant T (MAIT) cells are predominantly located in barrier tissues where they rapidly respond to pathogens and commensals by recognizing microbial derivatives of riboflavin synthesis. Early-life exposure to these metabolites imprints the abundance of MAIT cells within tissues, so we hypothesized that antibiotic use during this period may abrogate their development. We identified antibiotics that deplete riboflavin-synthesizing commensals and revealed an early period of susceptibility during which antibiotic administration impaired MAIT cell development. The reduction in MAIT cell abundance rendered mice more susceptible to pneumonia, while MAIT cell-deficient mice were unaffected by early-life antibiotics. Concomitant administration of a riboflavin-synthesizing commensal during antibiotic treatment was sufficient to restore MAIT cell development and immunity. Our work demonstrates that transient depletion of riboflavin-synthesizing commensals in early life can adversely affect responses to subsequent infections.

## Introduction

In a multicenter pediatric study, 73% of children received at least one course of antibiotics during their first year of life.^1^ While antibiotics can be beneficial, 26% of children across 32 hospitals were prescribed an antibiotic that was suboptimal due to ineffectiveness for the target bacteria, extended prophylaxis, or overly broad treatment.^2^ The widespread and often inappropriate use of antibiotics during infancy is associated with deleterious health outcomes later in life, including asthma and recurrent respiratory tract infections.^3–8^ Although antibiotics have been suggested to deplete commensal microbes that promote respiratory immunity,^9,10^ the mechanisms by which this occurs have not been established, preventing the development of effective therapeutics.

While conventional T cells recognize peptides presented by polymorphic major histocompatibility complex (MHC) proteins, many innate-like T cells are specific for non-peptidic microbial antigens, including lipids and metabolites, presented by monomorphic MHC class Ib molecules.^11^ Mucosal-associated invariant T (MAIT) cells comprise the predominant subset of innate-like T cells in most human tissues,^12^ reaching up to 9% of pulmonary T cells in healthy individuals.^13^ MAIT cells express semi-invariant T cell receptors (Vα19-Jα33 in mice and Vα7.2- Jα33/20/12 in humans) that recognize microbial derivatives of riboflavin (vitamin B2) synthesis presented by the MHC class-I related (MR1) molecule.^12,14^ Because MR1 is highly conserved between humans and mice,^15^ MAIT cells from both species respond to the same microbial metabolites. Riboflavin synthesis is broadly conserved among bacteria and fungi, so MAIT cells produce IFN-γ and/or IL-17A in response to a wide array of respiratory pathogens, with protection in murine models demonstrated for *Francisella tularensis, Klebsiella pneumoniae*, and *Legionella longbeachae*.^13,16–23^ While viruses do not synthesize metabolites, MAIT cells express receptors for the cytokines IL-12, IL-15, and IL-18, enabling them to release IFN-γ and granzyme B in response to viral infections,^24,25^ with protection against influenza shown in mice.^26^ The importance of these MR1-restricted T cells has been demonstrated in humans, where a single-nucleotide polymorphism in the *Mr1* gene is associated with increased susceptibility to tuberculosis,^27^ while a patient with recurrent bacterial and viral infections was found to have a nonfunctional MR1 variant.^28^ In children with bacterial or viral pneumonia, 30% of pulmonary MAIT cells produced IL- 17A compared to only 2% of conventional CD4^+^ T cells,^29^ demonstrating their outsized contribution to respiratory immunity.

Commensal microbes promote the development of MAIT cells through their synthesis of riboflavin intermediates.^30,31^ Although MAIT cells are nearly absent from the peripheral tissues of germ-free (GF) mice,^30–33^ microbial colonization of week-old GF neonates, but not 7-week-old GF adults, is sufficient to restore MAIT cell abundance.^30^ Since MAIT cells require early-life exposure to riboflavin-synthesizing commensals,^34^ we hypothesized that antibiotic use during this period may impede their development and subsequently impair respiratory immunity. Here we demonstrate that riboflavin-synthesizing bacteria are transiently enriched in early life and antibiotic treatment during this period is sufficient to deplete these commensals and hinder MAIT cell development. The resulting decrease in MAIT cell abundance renders adult animals more susceptible to bacterial pneumonia. Administering a riboflavin-synthesizing commensal during antibiotic treatment restores MAIT cell development and respiratory immunity, suggesting that probiotics may counter the detrimental effects of antibiotic use in early life.

## Results

### Identifying antibiotics that deplete riboflavin-synthesizing bacteria

While antibiotics alter the composition of the intestinal microbiome,^35^ the extent that they deplete riboflavin-synthesizing commensals has not been established. To identify antibiotics that may inhibit MAIT cell development, we cultured murine intestinal commensals with each of the 32 most prescribed antibiotics in neonatal intensive care units.^36^ The introduction of solid food during infancy causes diversification of the intestinal microbiome,^35^ so we isolated the cecal and colonic contents of wild-type (WT) C57BL/6J mice during weaning. Microbial samples were cultured anaerobically for 24 hours in chopped meat media supplemented with carbohydrates, which supports growth of a broad range of anaerobic bacteria,^37^ and the maximum mg/kg infant dose of each antibiotic as recommended by the U.S. Food and Drug Administration.^38^ Optical density measurements revealed that microbial growth was most strongly inhibited by β-lactam antibiotics, including ampicillin, piperacillin, cefazolin, amoxicillin, cefotaxime, and dicloxacillin (**Fig. 1A**).

**Figure 1.**
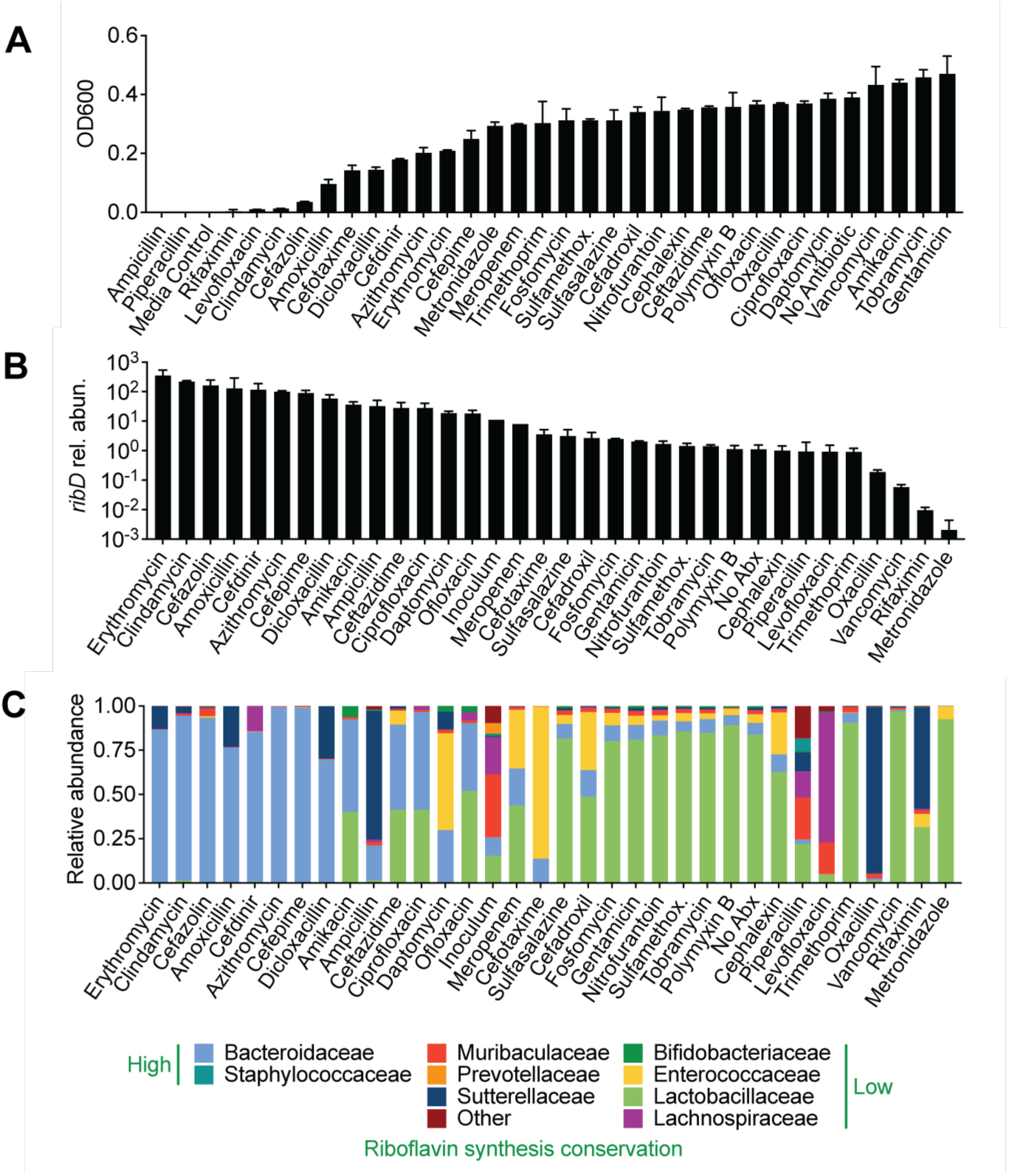
Identifying antibiotics that deplete riboflavin-synthesizing bacteria. *In vitro* assay of weaning-aged murine cecum and colon microbiota harvested anaerobically and cultured in rich media overnight with antibiotics. (A) Optical density (OD600) of microbial communities normalized to media controls after 24 hours of growth. (B) qPCR relative abundance of genomes with riboflavin synthesis gene *ribD* relative to 16Sv3v4, calculated using the Pfaffl method relative to the mean of no-antibiotic control samples. (A-B) Data represent the mean of duplicate samples ±SEM. (C) Family level 16S rRNA gene sequencing of microbial communities after 24h incubation with antibiotics. Data represent the mean of duplicate samples.

The *ribD* gene is necessary to generate the riboflavin derivative recognized by MAIT cells and indicative of the other riboflavin synthesis enzymes due to their organization within the riboflavin operon.^30,39,40^ To quantitate how antibiotics affect the riboflavin synthesis capacity of the intestinal microbiota, we designed degenerate primers that were complementary to highly conserved sequences within the *ribD* gene (**Supp. Fig. 1A**). After verifying that the assay was specific to *ribD* (**Supp. Fig. 1B**), we assessed the relative abundance of riboflavin synthesizers by quantitative polymerase chain reaction (qPCR). Although metronidazole and vancomycin had minimal effects on culture growth (**Fig. 1A**), they reduced the relative abundance of *ribD* by 493 and 17-fold relative to the untreated control respectively (**Fig. 1B**), suggesting outgrowth of non-riboflavin synthesizers. 16*S* rRNA gene sequencing revealed that both antibiotics allowed the growth of bacteria within the *Enterococcaceae* and *Lactobacillaceae* families (**Fig. 1C**), which exhibit low conservation of riboflavin synthesis among their species.^30^ Conversely, the relative abundance of *ribD* was unaffected or increased in cultures that were dominated by *Bacteroidaceae* (**Fig. 1C**), indicating growth of riboflavin-synthesizing species from this family. We concluded that antibiotics could reduce the relative abundance of riboflavin synthesizers, either by selective inhibition (e.g., vancomycin and metronidazole) or broad suppression of intestinal commensals (e.g., ampicillin).

### MAIT cells develop in response to bloom of riboflavin synthesizers during weaning

Oral β-lactam antibiotics are the first-line pediatric therapy for otitis media, pharyngitis, sinusitis, and urinary tract infections, while oral vancomycin and metronidazole are recommended for gastrointestinal infections.^41,42^ Consequently, ampicillin, vancomycin, and metronidazole are the 1^st^, 5^th^, and 44^th^ most administered drugs in neonatal intensive care units, respectively.^36^ Treatment courses are typically 10-14 days,^41,42^ so we determined whether this duration was sufficient to impair MAIT cell development *in vivo*. Mice received ampicillin for 2-week intervals followed by a fecal microbiome transplant (FMT) from untreated age-matched animals to limit the resulting microbial dysbiosis to the treatment period (**Fig. 2A**). Because the frequency of MAIT cells is highest in murine skin and correlates with their abundance in other tissues,^30^ we quantified cutaneous MAIT cells in adult 8-week-old animals. Ampicillin treatment during weeks 2-4 decreased MAIT cells by 2-3-fold, while treatment during weeks 1-3 and 3-5 had less severe effects (**Fig. 2B-D**). This period of susceptibility corresponded to the reported predominance of immature CD44^−^ MAIT cells within the thymi of 2-week-old mice, where they remain enriched through 4 weeks of age.^33^ Since mice begin weaning by 2 weeks of age, we hypothesized that the transition from nursing to solid food and the accompanying reduction of maternal immunoglobulins increases the abundance of riboflavin synthesizers within the intestinal microbiota. Longitudinal analysis of untreated mice revealed a predominance of *Lactobacillaceae* in the feces at 1 week, which resulted in a low capacity to synthesize riboflavin (**Fig. 2E-F**). The abundance of riboflavin synthesizers markedly increased by 2 weeks due to the emergence of *Bacteroidaceae* and *Tannerellaceae*, which have a high prevalence of this biosynthetic pathway.^30^ However, the accumulation of *Lachnospiraceae* and *Muribaculaceae* decreased the proportion of riboflavin synthesizers, resulting in a transient enrichment during weeks 2-4 (**Fig. 2E-F**).

**Figure 2.**
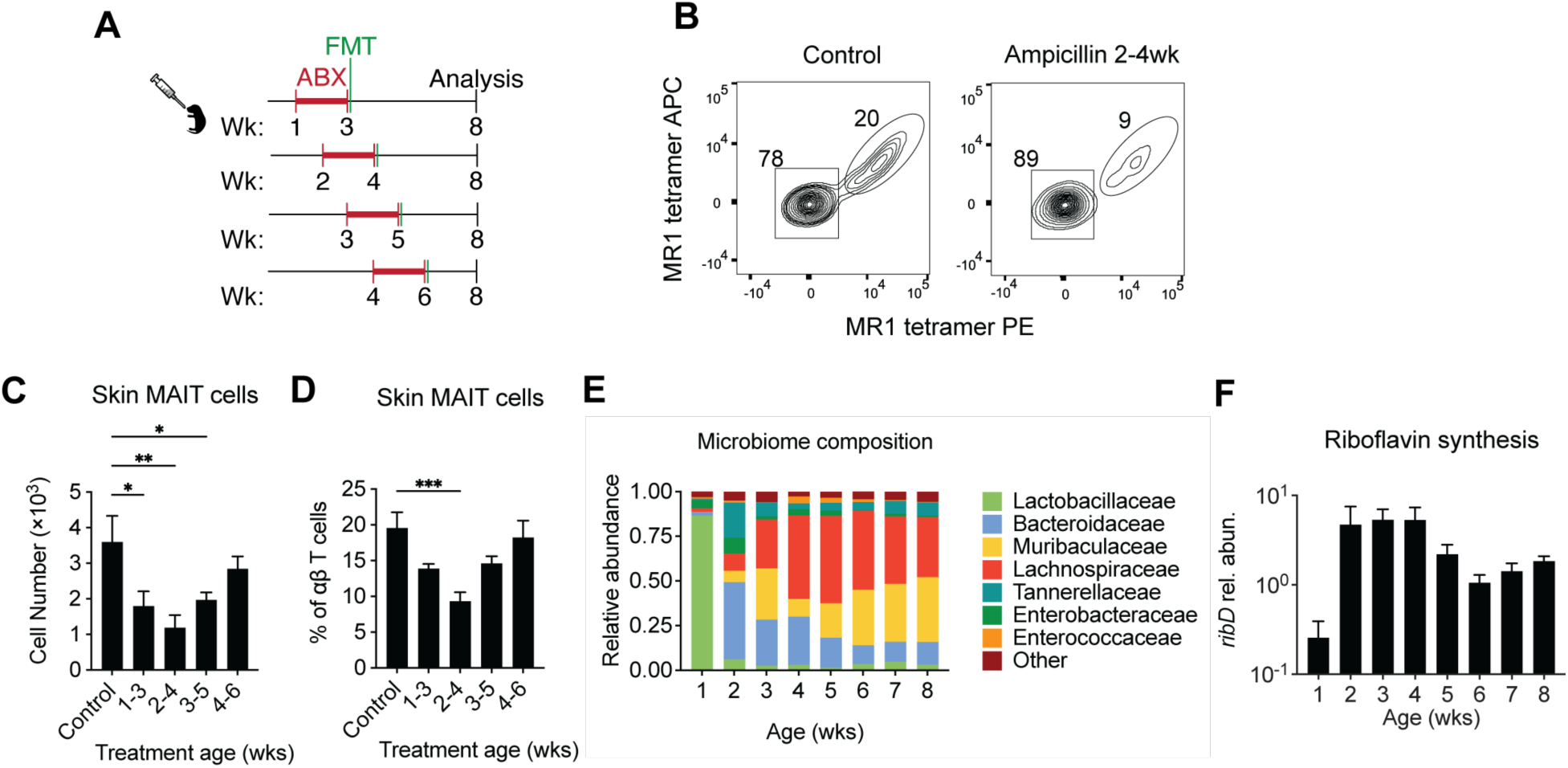
MAIT cells develop in response to bloom of riboflavin synthesizers during weaning. (A) Experimental schematic. (B) Multiparametric flow cytometry plots showing ear skin MAIT cell staining using dual MR1 tetramers loaded with 5-OP-RU. (C-D) MAIT cell number and frequency of αβ T cells. Data represent the mean ±SEM (n=5 per group). \**P* < 0.05, \*\**P* < 0.01, and \*\*\**P* < 0.001 as calculated with ordinary one-way ANOVA with Šidák’s multiple comparison correction. (E) 16*S* rRNA gene sequencing of microbiota collected weekly from untreated mice. Data represent the mean (n=3 mice per group). (F) qPCR of (E) showing relative abundance of genomes with riboflavin synthesis gene *ribD* relative to 16Sv3v4, calculated using the Pfaffl method relative to 6-week mean. Data represent the mean ±SEM (n=3 mice per group).

### Antibiotic use in early life preferentially inhibits MAIT cell development

Since a single course of ampicillin was sufficient to impair MAIT cell development, we assessed whether other T cell subsets were impacted by antibiotic use in early life. Mice received ampicillin from weeks 2-4 and were subsequently co-housed with untreated age-matched animals for a week to model cessation of antibiotics (**Fig. 3A**). Early-life ampicillin treatment had minimal effects on conventional CD4^+^ and CD8^+^ T cells and invariant natural killer T (iNKT) cells, but we observed a ∼1/3 reduction in γο T cells and regulatory T (T_reg_) cells within the skin and lungs (**Fig. 3B-C**). Prior work has demonstrated that Vγ6^+^ Vδ1^+^ T cells are reduced immediately following antibiotic treatment,^43^ but the persistence of this decrease weeks after microbial reconstitution has not been described. T_reg_ cells accumulate in murine skin during infancy in response to microbially-induced CCL20 and are reduced in GF animals.^44^ Since the accumulation of cutaneous T_reg_ cells peaks at day 13,^45^ the reduction caused by 2-4-week ampicillin treatment suggests that continuous microbial exposure is necessary to retain these cells within tissues. While early-life ampicillin treatment caused slight reductions of γο T cells and T_reg_ cells, MAIT cells were reduced by ∼3-fold across multiple tissues (**Fig. 3B-C**).

**Figure 3.**
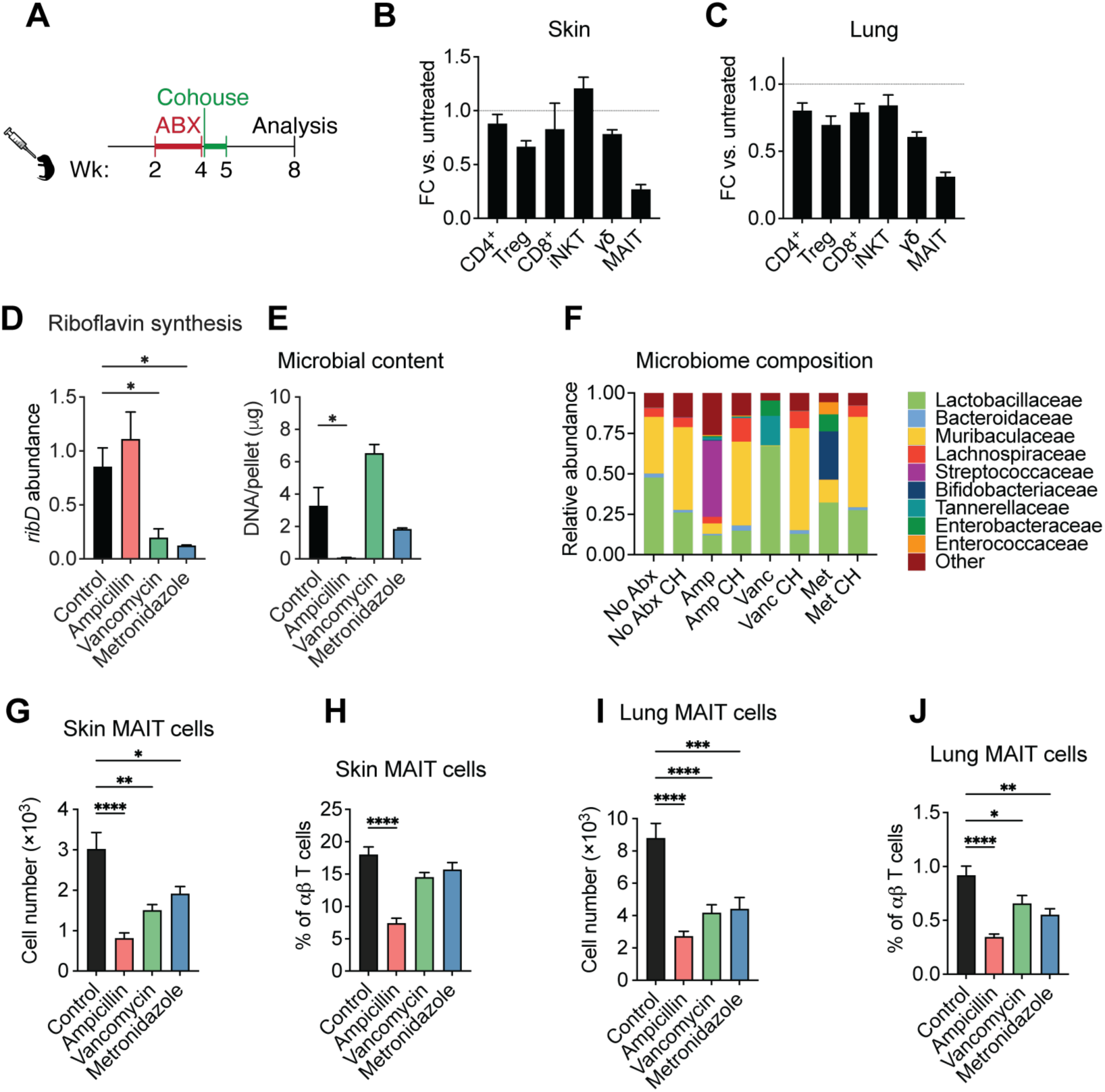
Antibiotic use in early life preferentially inhibits MAIT cell development. (A) Experimental schematic. (B-C) Lymphocytes fold change (FC) of adult mice following ampicillin treatment compared to untreated controls. Data represent the mean of three independent experiments (n=11-12 per group). (D) Relative abundance of genomes with riboflavin synthesis gene *ribD* relative to 16Sv3v4 after antibiotic treatment, calculated using the Pfaffl method relative to the mean untreated control samples. (E) Fecal microbial density after antibiotic treatment as determined by microbial DNA per fecal pellet. (D-E) Data represent the mean ±SEM (n=3 mice per group). \**P* < 0.05, as calculated with ordinary one-way ANOVA with Šidák’s multiple comparison correction. (F) 16S rRNA gene sequencing of microbiota from untreated mice and ampicillin- (Amp), vancomycin- (Vanc) or metronidazole- (Met) treated mice before and after co- housing (CH) with untreated mice for 1 week. Data represent the mean (n=3 mice per group). (G-J) MAIT cell number and frequency of αβ T cells after antibiotic treatment or control. \**P* < 0.05, \*\**P* < 0.01, \*\*\**P* < 0.001, and \*\*\*\**P* < 0.0001 as calculated with ordinary one-way ANOVA with Šidák’s multiple comparison correction. Data represent the mean of three independent experiments (n=10-12 per group).

Because our *in vitro* assay demonstrated that ampicillin broadly suppresses growth of intestinal commensals (**Fig. 1A**), the inhibition of MAIT cell development may have been due to the combined loss of microbial ligands for pattern recognition receptors and derivatives of riboflavin synthesis. To establish whether antibiotics that selectively deplete riboflavin synthesizers impair MAIT cell development, we administered vancomycin or metronidazole during the 2-4-week period. Feces collected at the conclusion of the antibiotic treatment revealed that vancomycin and metronidazole depleted the riboflavin synthesis capacity of the microbiota *in vivo* without inhibiting microbial growth (**Fig. 3D-E**), as observed *in vitro*. Both antibiotics permitted growth of bacterial families with a low conservation of riboflavin synthesis, including *Bifidobacteriaceae*, *Enterococcaceae*, and *Lactobacillaceae* (**Fig. 3F).** 2-4-week treatment with either vancomycin or metronidazole was sufficient to inhibit MAIT cell accumulation within the lungs and, to a lesser extent, the skin (**Fig. 3G-J**). Together, these results indicate that multiple frequently prescribed antibiotics can impair MAIT cell development when administered in early life.

### Early-life antibiotics impair immunity to bacterial pneumonia

Antibiotic use during the first 6 months of life increases the risk of recurrent respiratory tract infections in children.^3^ Since MAIT cells produce IFN-γ and/or IL-17A in response to a wide array of respiratory pathogens,^13,16–23^ we sought to determine if early-life antibiotic treatment could subsequently impair immunity to bacterial pneumonia.

The Gram-negative, facultative intracellular bacterium *F. tularensis* causes hundreds of infections each year, resulting in pneumonic tularemia when inhaled.^46^ Following retropharyngeal inoculation with the *F. tularensis* live vaccine strain (LVS), the abundance of MAIT cells within murine lungs increased more than 75-fold, while other T cell subsets did not proliferate extensively (**Fig. 4A**), which is consistent with prior studies.^20,21^ To establish the susceptibility of our mice, we inoculated WT animals with varying doses of *F. tularensis* LVS and monitored disease severity by weight loss and survival. Mice that received 1 × 10^3^ or more colony-forming units (CFUs) succumbed to the infection, while animals that received 6 × 10^2^ CFUs or less survived (**Fig. 4B-C**), indicating that the median lethal dose (LD_50_) is 6-10 × 10^2^ CFUs for our inoculation route.

**Figure 4.**
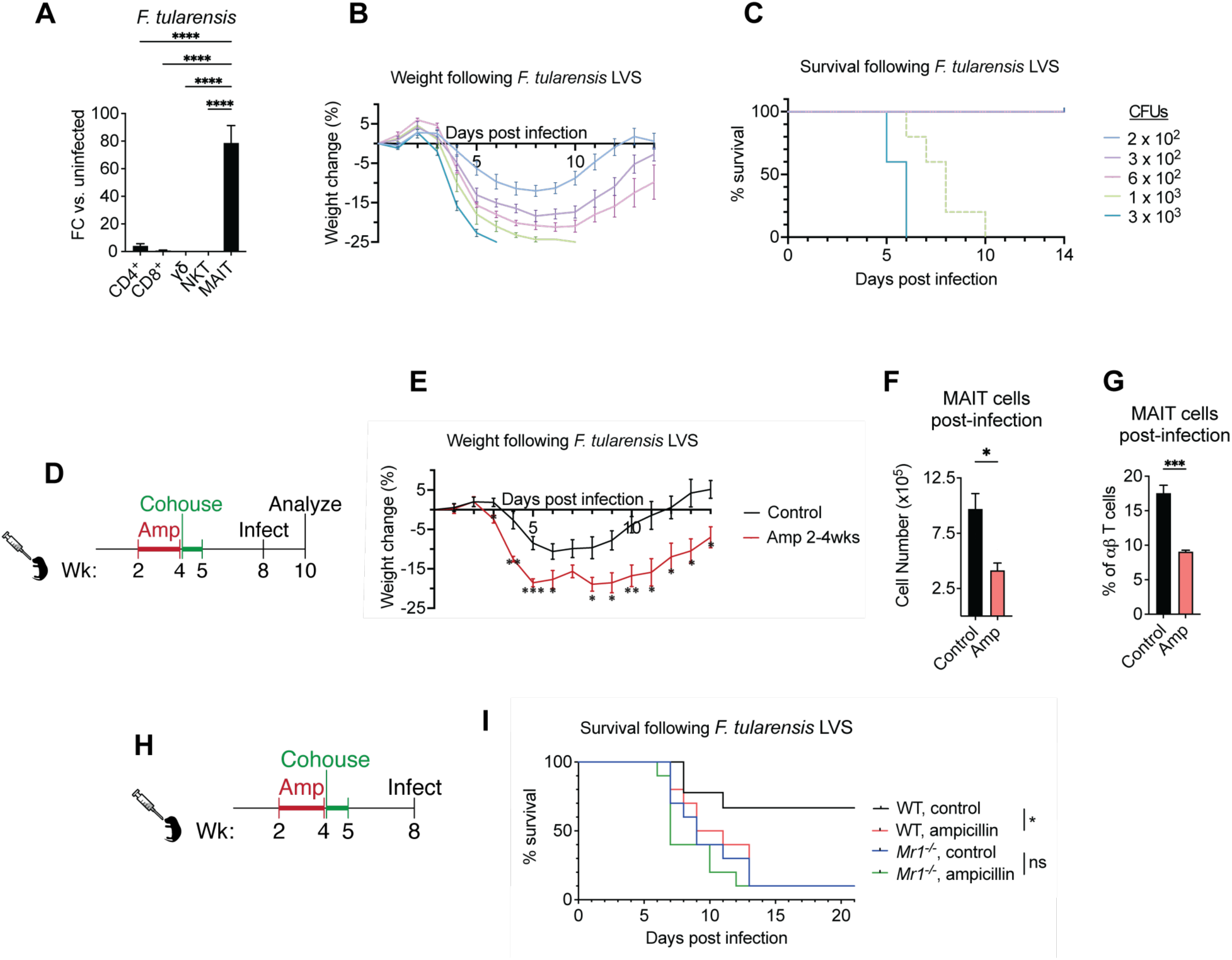
Early-life antibiotics impair immunity to bacterial pneumonia. (A) Fold change of lung lymphocytes after retropharyngeal inoculation with *F. tularensis* LVS. Data represents the mean ±SEM (n=3 per group) (B) Retropharyngeal dose titration morbidity by percent body weight. Data represent the mean ±SEM (n=5 per group). (C) Retropharyngeal dose titration mortality (n=5 mice per group). (D) Experimental schematic of early life antibiotic treatment followed by adult challenge with sublethal retropharyngeal *F. tularensis* LVS dose. (E) Weight loss of ampicillin-treated or control mice following sublethal retropharyngeal *F. tularensis* LVS dose. Data represent the mean ±SEM (n=4-5 per group). \**P* < 0.05, \*\**P* < 0.01, \*\*\**P* < 0.001 using Student’s t-test. (F-G) Lung MAIT cell number and frequency of αβ T cells 14 days post- infection with sublethal retropharyngeal *F. tularensis* LVS dose in (E). Data represent the mean ±SEM. \**P* < 0.05, and \*\*\**P* < 0.001, as calculated with ordinary one-way ANOVA. (H) Experimental schematic of early life antibiotic treatment followed by adult challenge with LD_50_ retropharyngeal *F. tularensis* LVS dose. (I) Mortality of ampicillin-treated or control mice following LD_50_ retropharyngeal *F. tularensis* LVS dose in wild type and *Mr1^-/-^* mice (n=9-10 per group). \**P* < 0.05 as calculated with Mantel-Cox test.

To establish whether early-life antibiotic use impairs pulmonary immunity, we inoculated ampicillin and vehicle control-treated mice with a sublethal 4 × 10^2^ CFU dose of *F. tularensis* LVS (**Fig. 4D**). Mice that received antibiotics in early life lost twice as much body weight and failed to recover 14 days post-infection (**Fig. 4E**). Ampicillin-treated mice had half the number and frequency of MAIT cells compared to control animals (**Fig. 4F-G**), indicating that the antibiotic-mediated impairment of MAIT cells persisted throughout the infection. Following an LD_50_ dose of 8 × 10^2^ CFUs of *F. tularensis* LVS (**Fig. 4H**), WT mice that received early-life ampicillin were significantly more susceptible than vehicle control-treated animals, with a survival rate that mirrored MAIT cell-deficient *Mr1^-/-^* mice (**Fig. 4I**). Ampicillin-treated *Mr1^-/-^* mice succumbed to the infection similarly to the control *Mr1^-/-^* animals, demonstrating that antibiotic-mediated impairment of immunity is mediated by the loss of MAIT cells.

### Probiotic restoration of MAIT cell-mediated immunity

Due to the abundance of *Bacteroidaceae* species during the 2-4-week period (**Fig. 2F**) and their susceptibility to vancomycin and metronidazole (**Fig. 1C**), we hypothesized that a riboflavin-synthesizing member of this family may be sufficient to restore MAIT cell development. We identified the human fecal isolate *Bacteroides thetaiotaomicron* strain VPI-5482, which retains the riboflavin synthesis pathway and the PER-1 β-lactamase that hydrolyzes penicillins and cephalosporins,^47^ enabling coadministration during ampicillin treatment. As a non-riboflavin synthesizing control, we selected an ampicillin-resistant *Enterococcus faecium* isolated from human feces. Mice received daily oral gavages of either *B. thetaiotaomicron* or *E. faecium* during the ampicillin treatment from 2-4 weeks of age (**Fig. 5A**). Concomitant administration of *B. thetaiotaomicron*, but not *E. faecium*, was sufficient to increase the number and frequency of pulmonary MAIT cells in adult mice (**Fig. 5B-C**). Mice that received *E. faecium* had a slight reduction in MAIT cells, which may be due to diminished colonization of riboflavin synthesizers during the co-housing period. To determine whether administration of a riboflavin-synthesizing probiotic is sufficient to restore immunity, mice were inoculated with an LD_50_ dose of 8 × 10^2^ CFUs of *F. tularensis* LVS at 8 weeks of age (**Fig. 5D**). Animals that received *B. thetaiotaomicron* were less susceptible than mice that received *E. faecium* or no probiotic during ampicillin treatment, with survival rates comparable to control mice that were not administered antibiotics (**Fig. 5E**). Thus, early-life antibiotic use subsequently impairs MAIT cell-mediated immunity by reducing riboflavin synthesis capacity, which can be ameliorated by administering a riboflavin-synthesizing probiotic.

**Figure 5.**
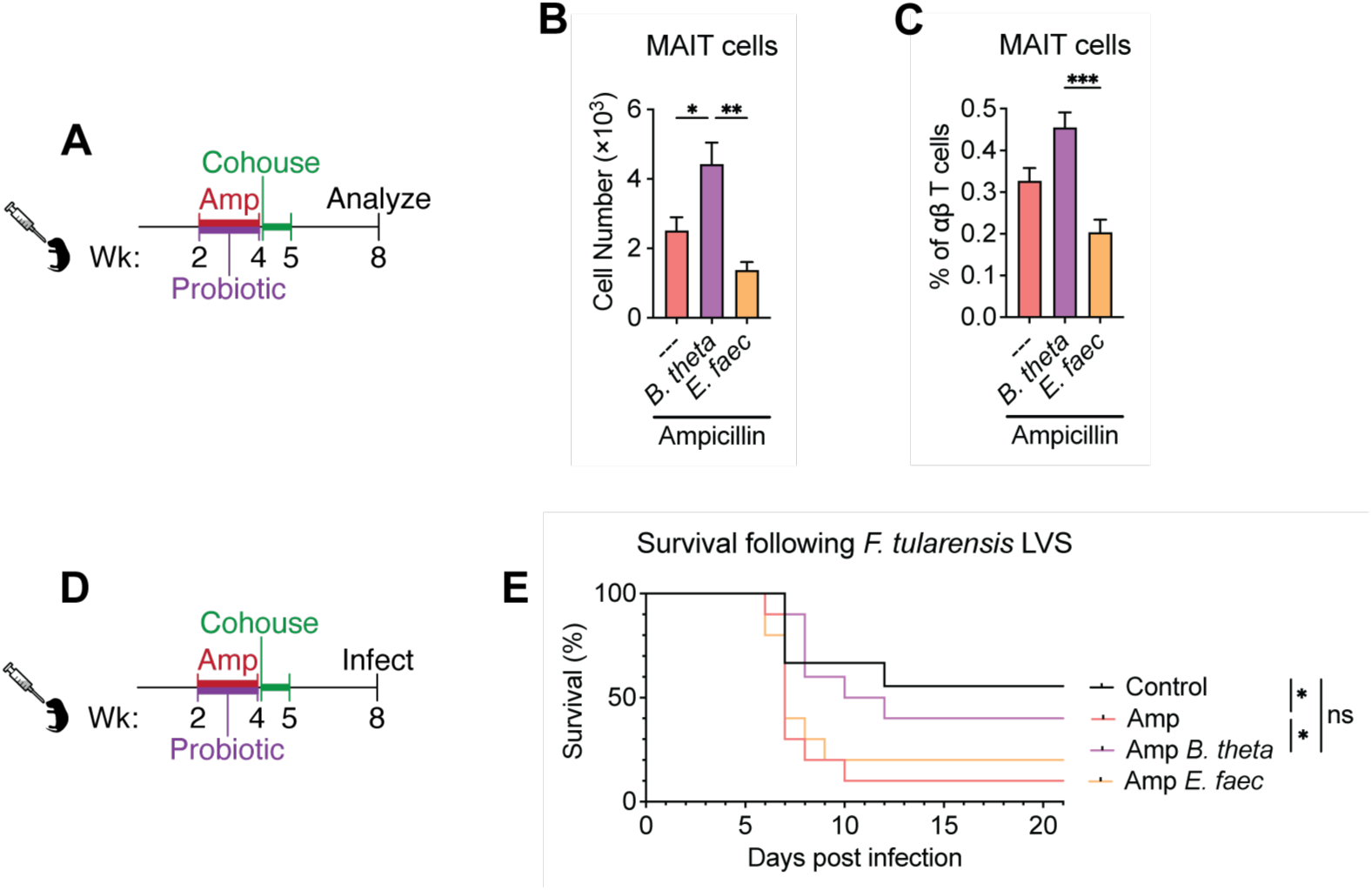
Probiotic restoration of MAIT cell-mediated immunity. (A) Experimental schematic of early life antibiotic treatment with concurrent probiotic administration. (B-C) Lung MAIT cell number and frequency of αβ T cells after antibiotic treatment with or without riboflavin-proficient *Bacteroides thetaiotaomicron* probiotic or riboflavin-deficient *Enterococcus faecium* probiotic. Data represent the mean of two independent experiments ±SEM (n=7-11 per group). \**P* < 0.05, \*\**P* < 0.01, \*\*\**P* < 0.001, and \*\*\*\**P* < 0.0001 as calculated with ordinary one-way ANOVA with Šidák’s multiple comparison correction. (D) Experimental schematic of early life antibiotic treatment with concurrent probiotic administration followed by adult challenge with LD_50_ retropharyngeal *F. tularensis* LVS dose. (E) Mortality of ampicillin-treated mice with or without riboflavin-proficient *Bacteroides thetaiotaomicron* probiotic or riboflavin- deficient *Enterococcus faecium* probiotic and control mice following LD_50_ retropharyngeal *F. tularensis* LVS dose (n=9-10). Statistics were performed using Mantel-Cox test.

## Discussion

Our findings reveal a transient enrichment of riboflavin-synthesizing intestinal commensals in early life that is necessary for the development of peripheral MAIT cells. Predictive functional analysis of human stool determined that the riboflavin biosynthesis pathway is more abundant in children than adults,^48^ so there may be a corresponding period for microbial induction of MAIT cells in humans. Within the first year of life, the frequency of MAIT cells in human blood increases approximately 10-fold and these cells acquire their capacity to produce cytokines.^49^ During this period, the infant intestinal microbiome is primarily composed of bacteria from the families *Enterobacteriaceae*, *Bacteroidaceae*, and *Clostridiaceae*,^35^ which have a high prevalence of riboflavin synthesis.^30^ However, their collective abundance decreases following the first year,^35^ suggesting a transient enrichment in riboflavin synthesizers during infancy. Although it remains unclear whether human MAIT cell development depends on this period, children that receive hematopoietic stem cell transplants regain only 10% of their pre-transplant MAIT cell frequencies a year later, even though conventional T cells fully recover.^50^

Our *in vitro* screen identified multiple antibiotics that depleted riboflavin synthesizers within the murine intestinal microbiota, including ampicillin, vancomycin, and metronidazole. Both ampicillin and vancomycin decreased microbial diversity within the human infant microbiota and early-life antibiotic treatment had persistent effects on the microbial composition.^51^ We found that mice treated with ampicillin, vancomycin, or metronidazole from 2-4 weeks of age developed fewer MAIT cells in their tissues and this impairment persisted into adulthood. The prevalence of immature CD161^−^ MAIT cells within the blood of human neonates and infants indicates that extrathymic signals are necessary for their development in early life.^33,49^ Given that the microbiota promotes maturation of murine MAIT cells,^30,31,33^ the development of human MAIT cells is likely vulnerable to antibiotic use during infancy. Disparate antibiotic use in early life may explain why the range of MAIT cell frequencies in pediatric and adult peripheral blood is broader than the abundance in neonatal cord blood.^49^

Following early-life antibiotic treatment, mice were more susceptible to bacterial pneumonia later in life. In addition to increasing the risk of recurrent respiratory tract infections in children,^3^ antibiotic use during infancy more than doubles the odds of developing pediatric asthma,^4–7^ with the risk increasing approximately 20% with each course of antibiotics.^4,8^ Since MAIT cells have been shown to restrict allergic airway inflammation in response to *Alternaria alternata* and house dust mite extracts,^52,53^ additional research is warranted to assess whether the increased prevalence of asthma is due to the depletion of MAIT cells. Additionally, the decrease in cutaneous MAIT cells following early-life antibiotics suggest that immunity within the skin may also be impaired. Antibiotic use during infancy has been shown to increase the risk of atopic dermatitis in children,^8^ although it remains unclear whether MAIT cells are beneficial or detrimental during this inflammatory skin disease.^54,55^

To maintain the riboflavin synthesis capacity of the microbiome during antibiotic treatment, we administered a human fecal isolate of *B. thetaiotaomicron*, which was sufficient to increase MAIT cell abundance and restore respiratory immunity. Due to the abundance of *Bacteroidaceae* within the intestinal microbiota of infants,^35^ members of this bacterial family may be essential for MAIT cell development. However, we demonstrated that multiple antibiotics depleted *Bacteroidaceae* from the murine microbiota. Human newborns that received the β-lactams amoxicillin and cefotaxime had less *Bacteroides* and the abundance of multiple taxa within this genus were decreased over a year following antibiotic treatment.^56^ Identifying which riboflavin synthesizers are the primary determinants of MAIT cell abundance and developing therapeutics to restore them is of interest for human health.

Our work identifies weaning as a critical period for the development of MAIT cells, which depend on the transient enrichment of riboflavin-synthesizing commensals in early life. Antibiotics that are readily prescribed to infants can deplete riboflavin synthesizers and administration during this period abrogates MAIT cell development, rendering adult mice more susceptible to infection. Concomitant administration of a riboflavin-synthesizing commensal during antibiotic treatment was sufficient to restore murine MAIT cell development and immunity, indicating that probiotic supplementation may ameliorate the immunologic consequences of early-life antibiotic use.

## Acknowledgements

The authors thank the Scripps Research Department of Animal Resources, Flow Cytometry Core, and Genomics Core. This work was supported by the National Institutes of Health (K22AI146217, R21AI171697, & R35GM151347 to M.G.C.), the National Science Foundation (Graduate Research Fellowship to G.L.), and Scripps Research.

## Author contributions

Conceptualization: A.L.S. and M.G.C.; Methodology: A.L.S., J.M., and M.G.C.; Investigation: A.L.S., J.M., D.H, G.L., and A.C.; Analysis: A.L.S. and M.G.C.; Writing – Original Draft: A.L.S. and M.G.C.; Writing – Review & Editing: A.L.S., J.M., D.H, G.L., and M.G.C.; Visualization: A.L.S. and M.G.C.; Supervision: M.G.C.; Funding Acquisition: M.G.C., J.M., and G.L.

## Declaration of interests

A.L.S. and M.G.C. are listed as inventors on a patent application describing probiotic restoration of MAIT cells. All other authors declare no competing interests.

## STAR Methods

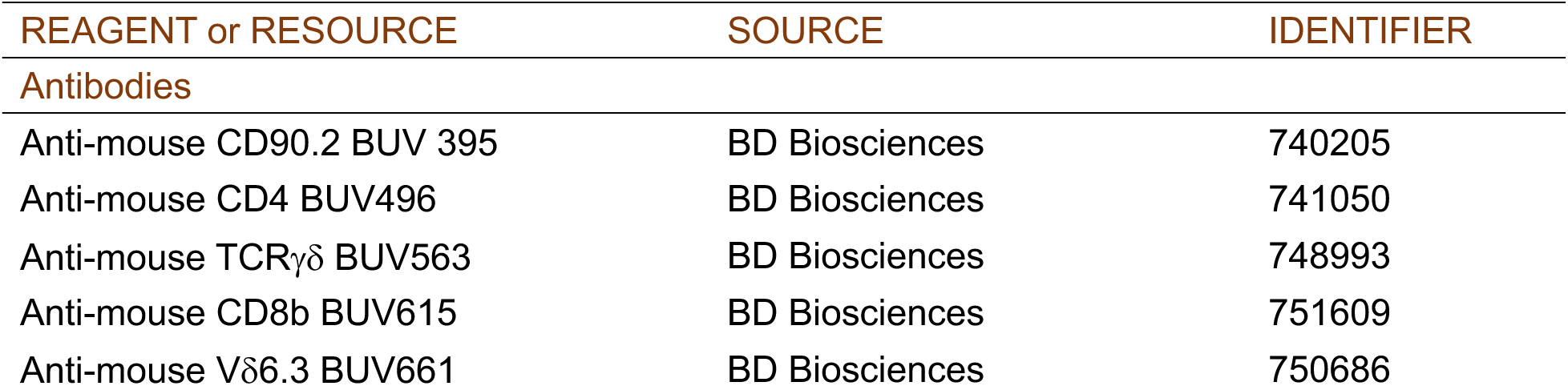
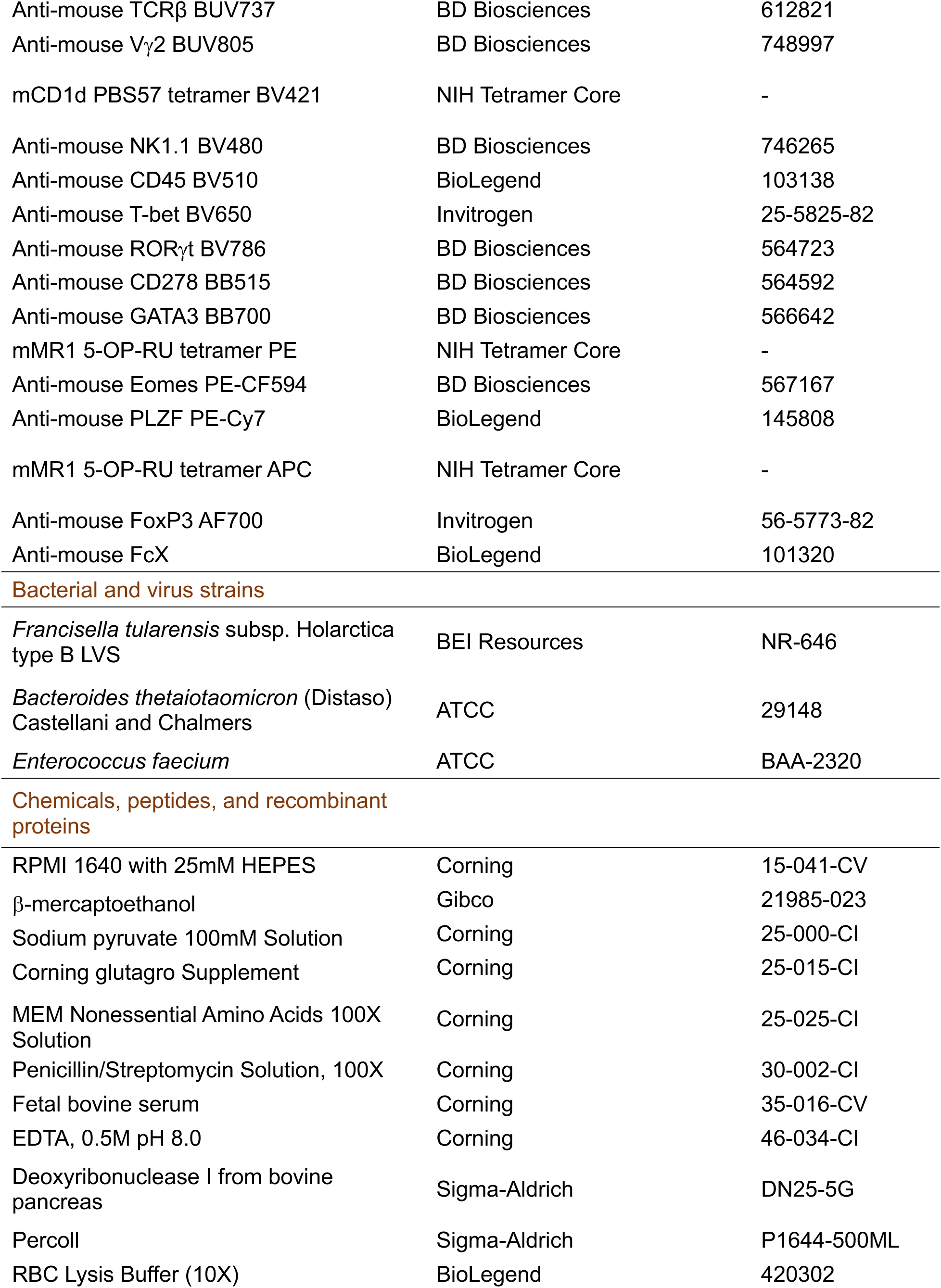
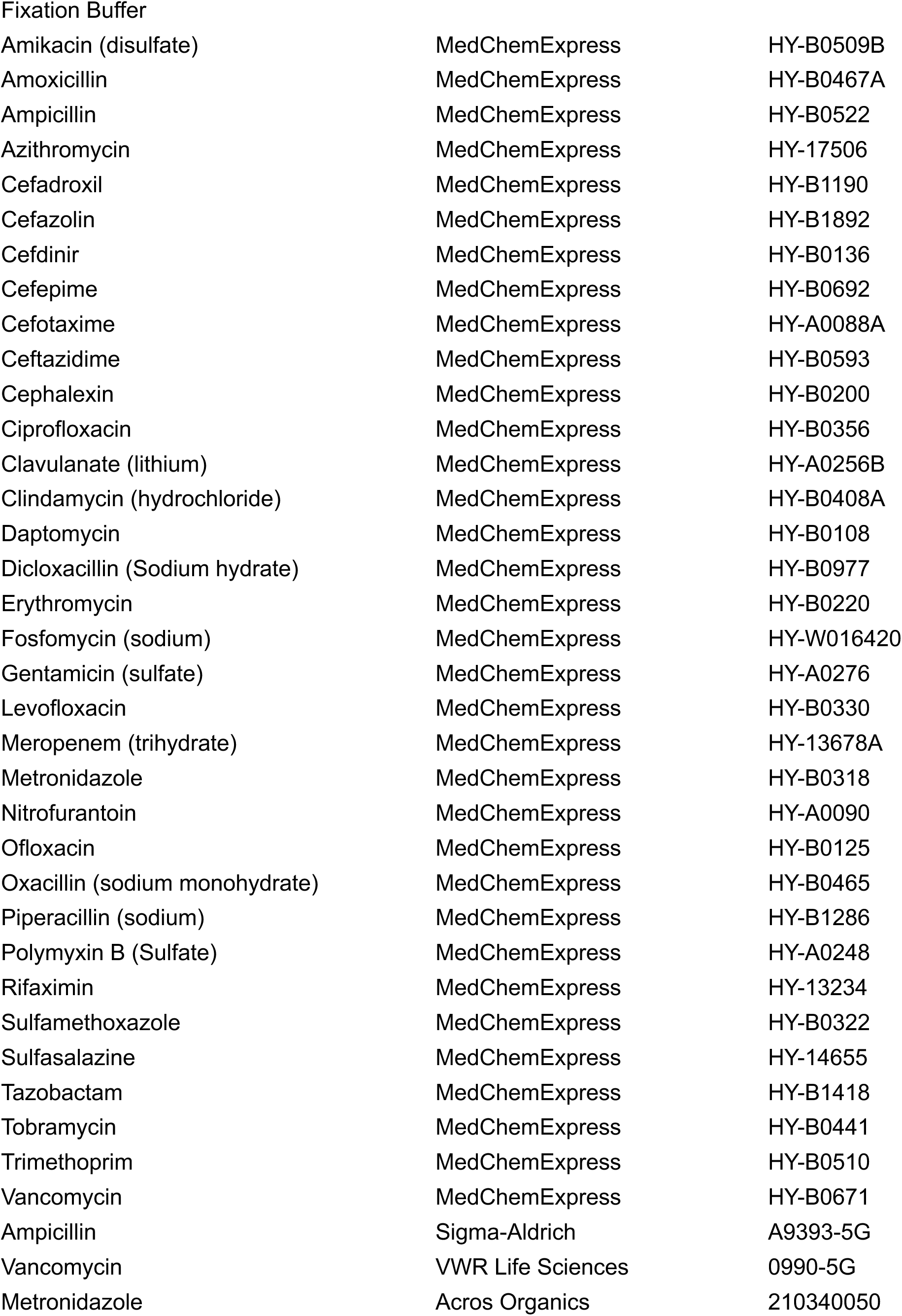
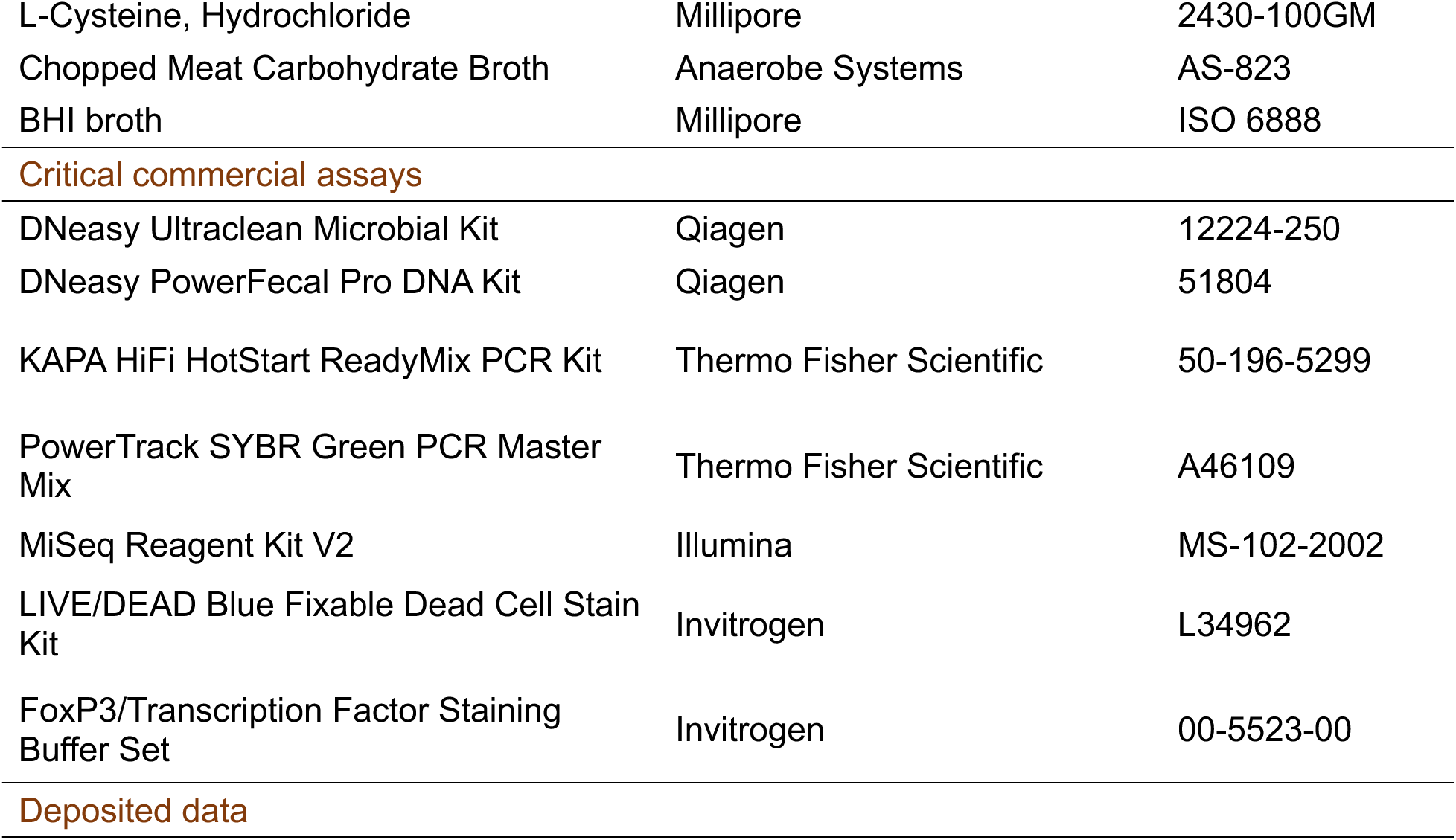
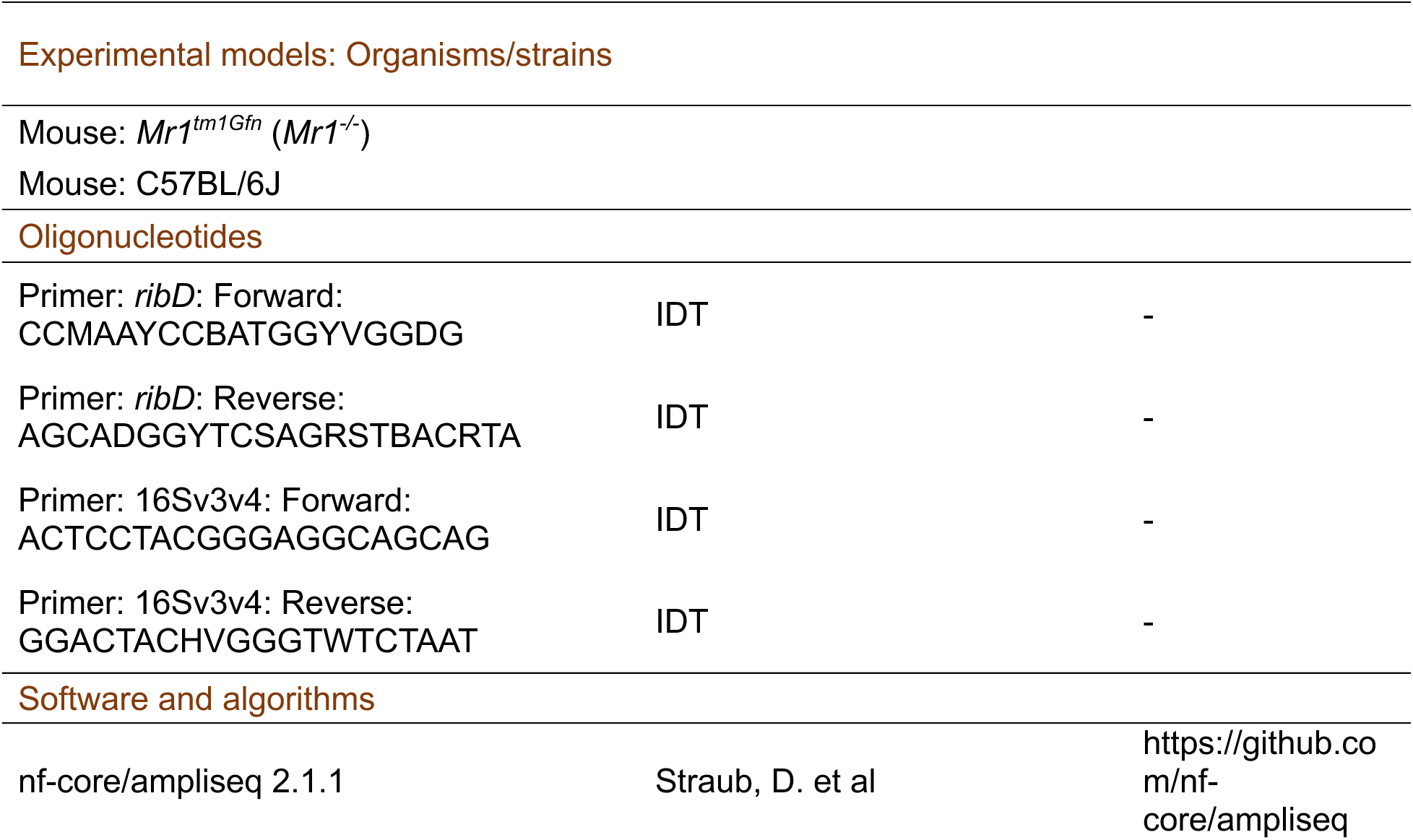
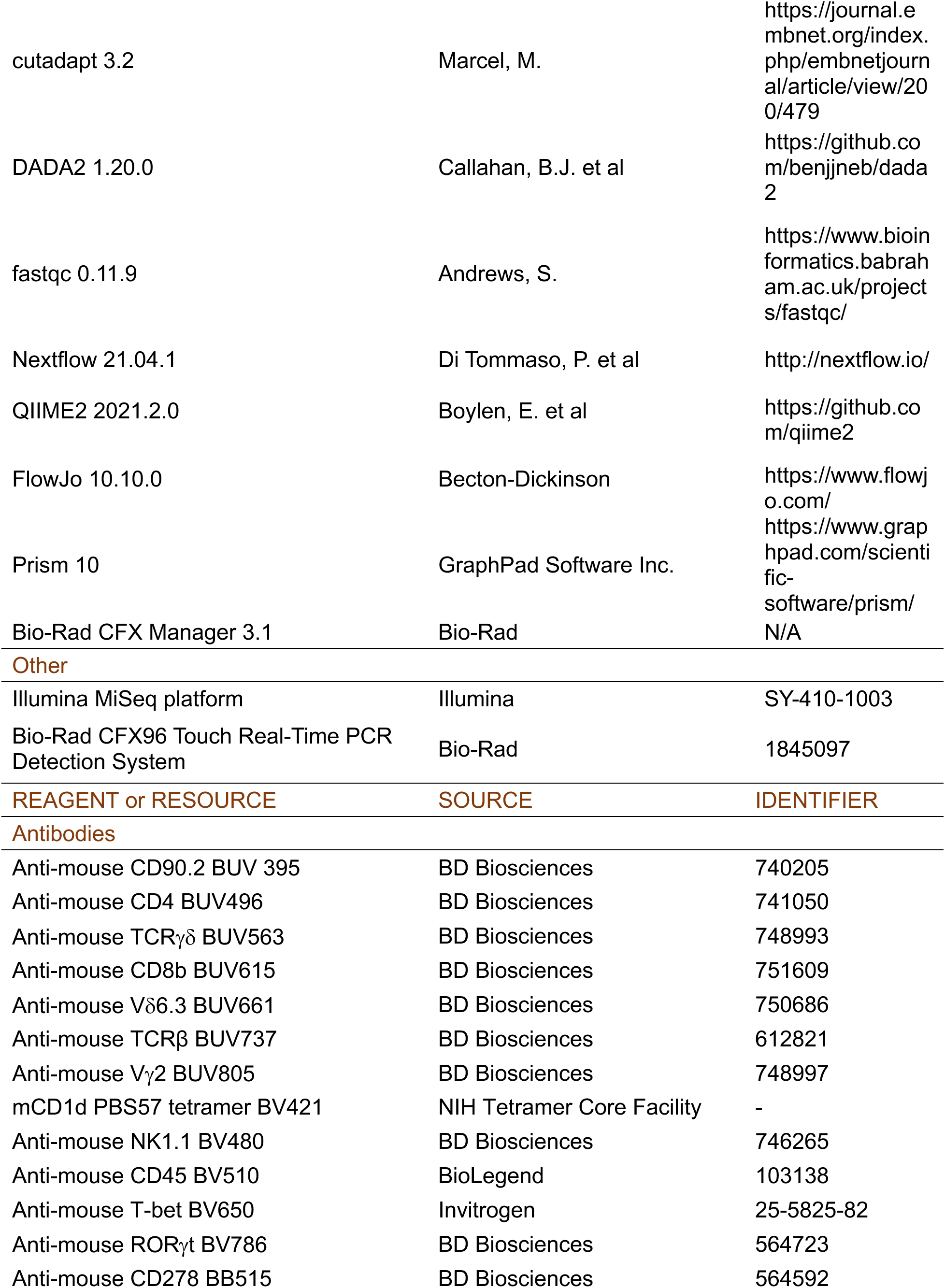
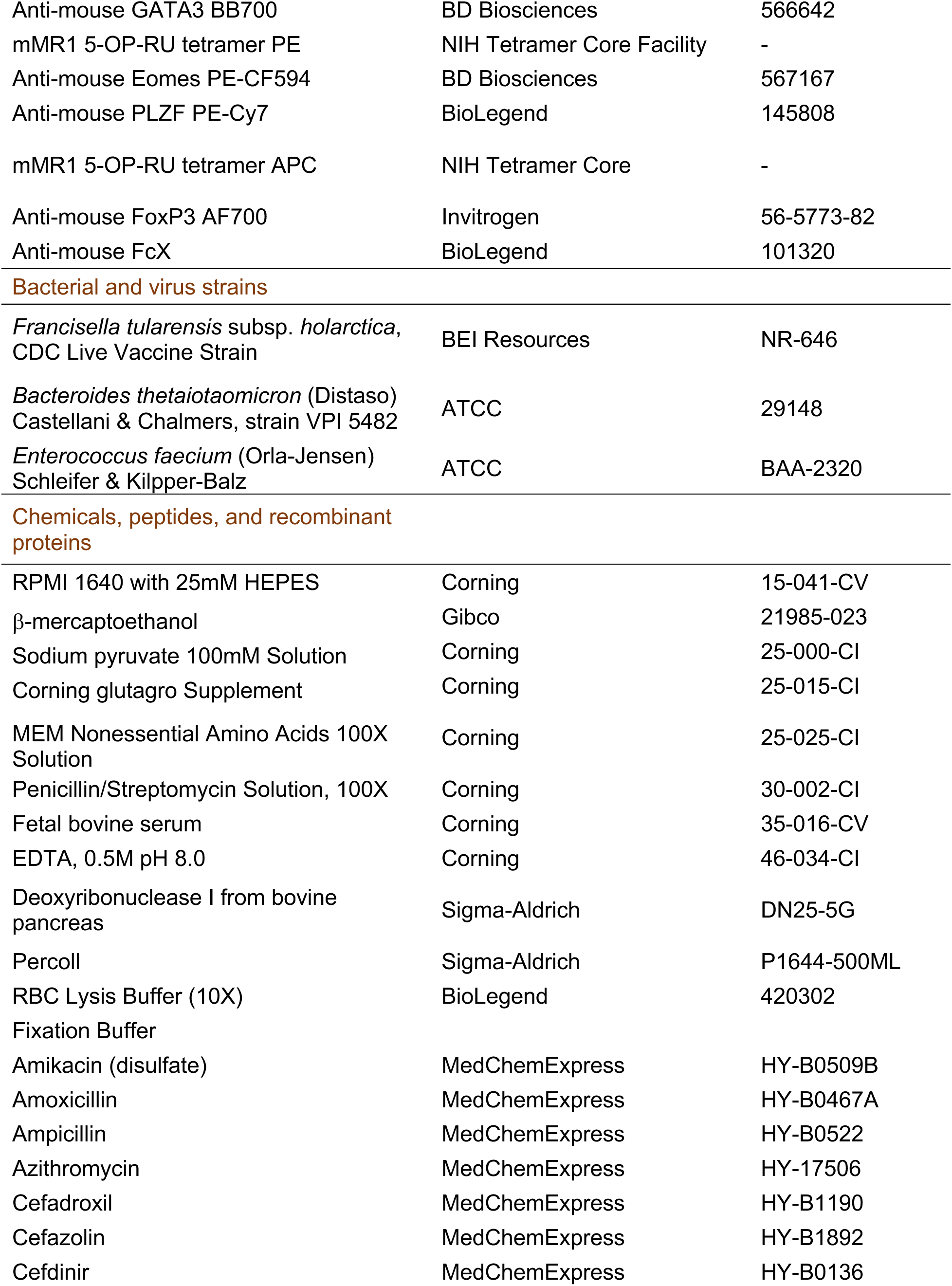
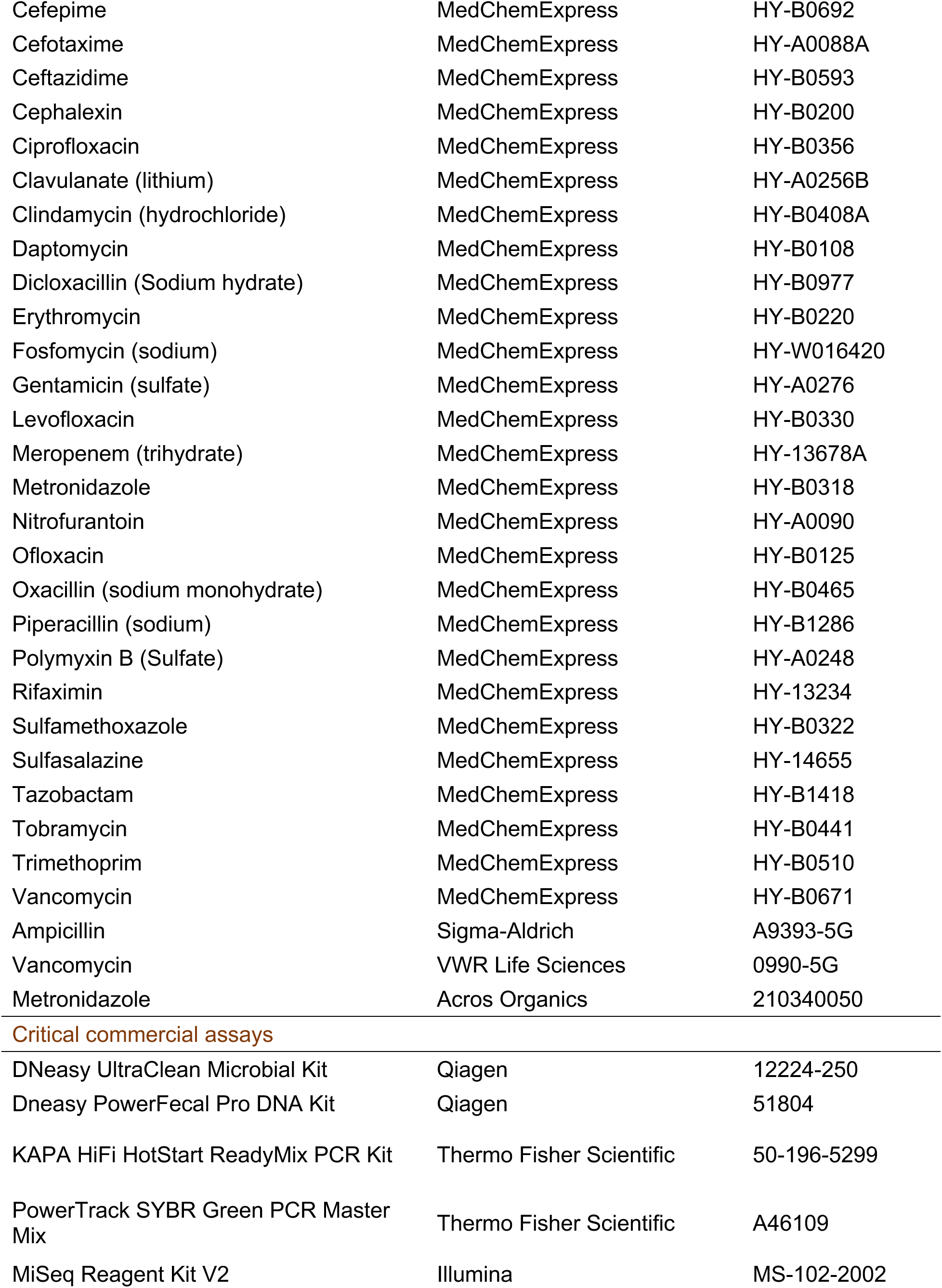
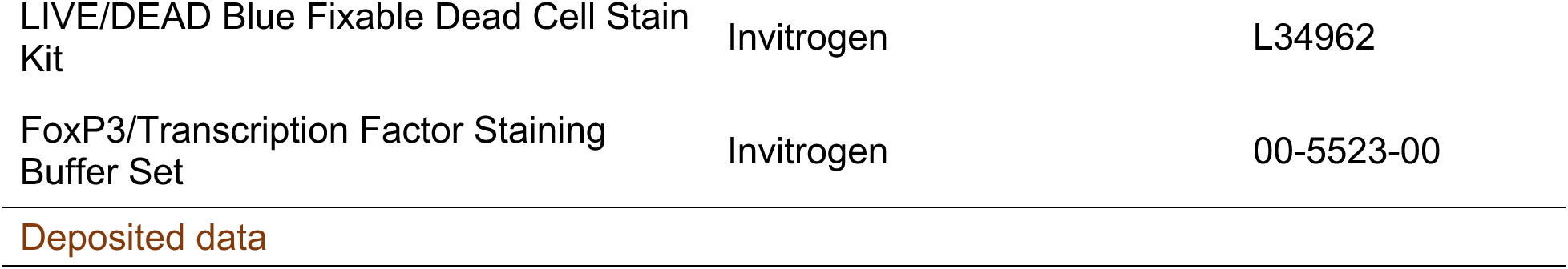
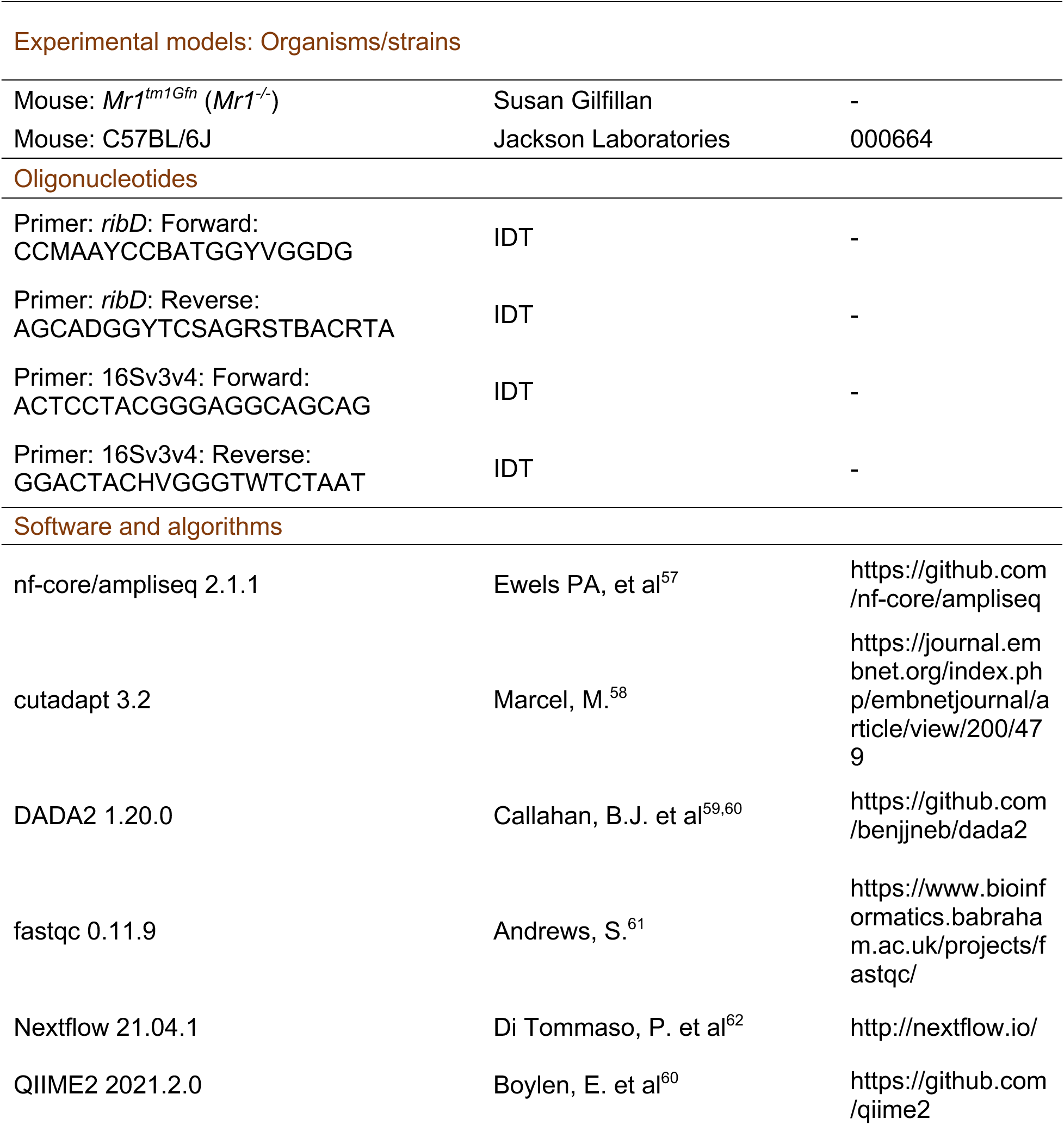
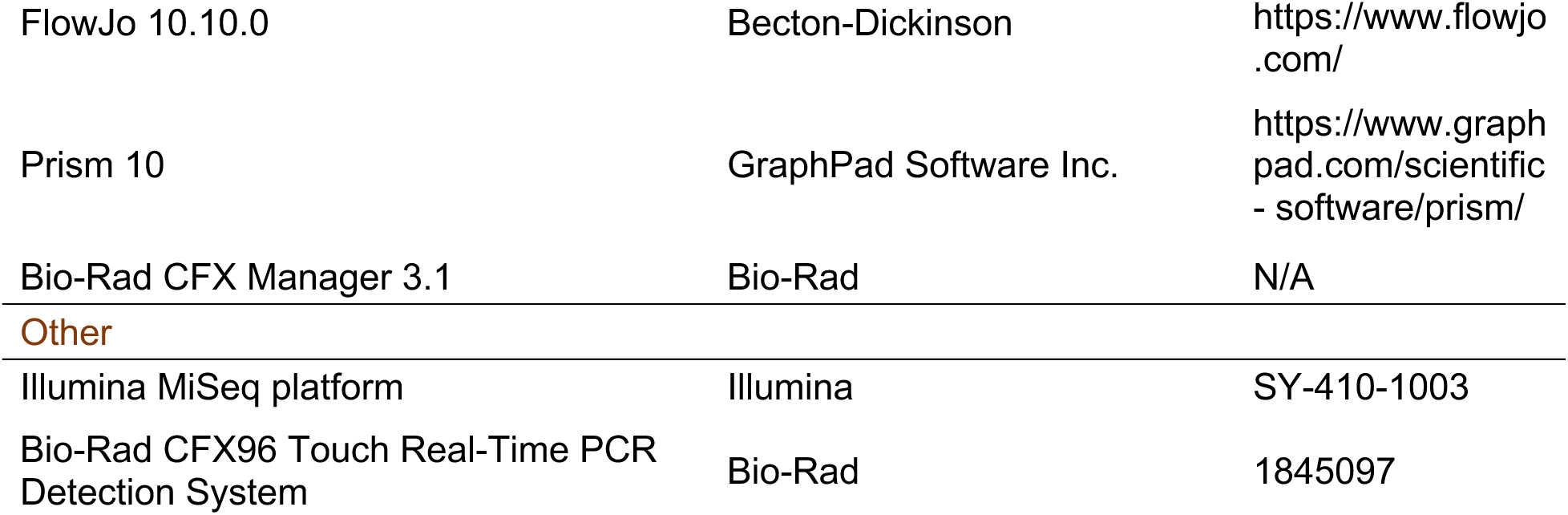
KEY RESOURCES TABLE.

## RESOURCE AVAILABILITY

### Lead Contact

Further information and requests for resources and reagents should be directed to and will be fulfilled by the lead contact, Michael G. Constantinides.

### Materials Availability

This study did not generate new unique reagents.

### Data and Code Availability

- Sequencing data will be deposited in a repository upon publication.
- All data reported in this paper will be shared by the lead contact upon request.
- This paper does not report original code.
- Any additional information required to reanalyze the data reported in this paper is available from the lead contact upon request

## EXPERIMENTAL MODELS AND SUBJECT DETAILS

### *In vitro* antibiotic screen

The antibiotic screen was adapted from previous work to analyze early life microbiota ^63^. Whole digestive tract from proximal small intestine to rectum were excised from 2 each of two-and-a-half and three-week-old mice and quickly transferred into an anaerobic chamber (Coy Lab Products). Cecal and colon contents were harvested, passed through a 70μm filter using 5 ml of prereduced PBS with 0.1% cysteine, diluted 12-fold with Chopped Meat Carbohydrate Broth (Anaerobe Systems), and adjusted to 0.1 OD after subtracting media blank. 0.8ml aliquots were distributed into deep-well 96-well plates in duplicate with each antibiotic. Antibiotics were acquired from MedChemExpress in 10mM DMSO or water. Antibiotic concentrations were derived from maximum neonatal or pediatric mg/kg dose^38^. This dose was adapted to a molar assay concentration by assuming homogenous diffusion and 70% water by mass. Assay concentration was limited to 150μM to limit DMSO concentration to 1.5%. DMSO concentration was normalized to all samples and controls in mother dilutions which were prepared with 25x concentration for transfer to the 96-well deep-well assay plate. Samples were incubated for 24h at 37°C. 100μl of each sample was transferred to an optically clear 96-well plate for optical density measurement. DNA was purified from the remainder using DNeasy Ultraclean Microbial Kit (Qiagen) for downstream analysis.

### *ribD* qPCR assay

qPCR analysis of riboflavin synthesis capacity was performed using PowerTrack SYBR Green (Applied Biosystems, A46109) with a Bio-Rad C1000 Touch Thermal Cycler with CFX96 Real-Time System. *RibD* primers were designed using a multiple alignment of reference genomes from 4 riboflavin-synthesizing bacterial species previously isolated from our facility: *Enterocloster bolteae*, *Bacteroides thetaiotaomicron*, *Muribaculum intestinale*, and *Duncaniella spp;* RibD-F (5’CCMAAYCCBATGGYVGGDG-3’; RibD-R: 5’AGCADGGYTCSAGRSTBACRTA-3’; B=C,G,T, D=A,G,T, M=A,C, R=A,G, S=C,G, V=A,C,G, Y=C,T). V3-V4 hypervariable region of the bacterial 16*S* rRNA gene was amplified with 338F and 806R primers (338F: 5′- ACTCCTACGGGAGGCAGCAG-3′; 806R: 5′-GGACTACHVGGGTWTCTAAT-3′). *ribD* relative abundance was quantified using the Pfaffl method relative to 16Sv3v4, relative to no-antibiotic control samples^64^.

### 16S rRNA gene sequencing from *in vitro* cultures

Amplification of the V3-V4 hypervariable region of the bacterial 16*S* rRNA gene was performed using 338F and 806R primers similar to those previously described^63^, which contained 12-base sample barcodes (N), unique molecular identifiers (n) to correct for PCR duplication artifacts, and Illumina sequencing components (5′- AATGATACGGCGACCACCGAGATCTACACTCTTTCCCTACACGACGCTCTTCCGATCT NNNNNNNNNNNNACTCCTACGGGAGGCAGCAG-3′) and reverse primer (5′-CAAGCAGAAG ACGGCATACGAGATNNNNNNNNNNNNGTGACTGGAGTTCAGACGTGTGCTCTTCCGATCTn nnnnnnnnnGGACTACHVGGGTWTCTAAT-3′). Amplification was performed using 50ng DNA, 10μl 2× KAPA HiFi HotStart ReadyMix (Kapa Biosystems), and 0.35μM of each primer. Samples with low DNA concentration were amplified using a maximum volume. PCR cycling was performed in 2 stages: 95 °C for 3 min, 5 cycles of 98°C for 20sec, 57°C for 15sec to optimize primer binding to genomic amplicon region, 72°C for 45sec, and a final 1min elongation at 72°C. This was followed by 95 °C for 3 min, 20 cycles of 98°C for 20sec, 67°C for 15sec to optimize primer binding to PCR amplicon, 72°C for 45sec, and a final 1min elongation at 72°C. Samples with abundant template concentration were pooled and purified using Agencourt AMPure XP beads (Beckman Coulter). Samples with low template concentration were purified individually and quantified using Quant-iT dsDNA Assay Kit (Thermo Fisher) prior to pooling. Sequencing was performed using Illumina MiSeq. Sequencing reads were demultiplexed using second barcode indices and processed to remove PCR duplication artifacts using unique molecular identifier sequences^63^. Sequences were analyzed for taxonomic identification using the nf-core ampliseq workflow with FastQC, Cutadapt, MultiQC, QIIME2, and DADA2.

### 16S rRNA gene sequencing from feces

Fecal DNA was purified using the QIAmp PowerFecal Pro DNA Kit (Qiagen), amplified and analyzed as previously described. Sequencing was performed using Illumina Nextseq.

### Mice

Wild-type (WT) C57BL/6J mice were acquired from the rodent breeding colony at Scripps Research, which is supplied by The Jackson Laboratory. *Mr1^tm1Gfn^* (*Mr1^-/-^*) mice were generated by Dr. Susan Gilfillan^32^. Mice were bred and cared for in a facility accredited by the American Association for the Accreditation of Laboratory Animal Care (AAALAC) at the Scripps Research Institute. All experiments were conducted at Scripps Research in accordance with an Animal Safety Protocol (21-0003) approved by the Scripps Research Institutional Animal Care and Use Committee (IACUC). All experiments were age- and sex-matched.

### Antibiotic treatment

*In vivo* antibiotic doses were adapted from human neonatal and pediatric doses at or below maximum daily mg/kg dose^38^; Ampicillin 100mg/kg/d, Vancomycin 40 mg/kg/d, Metronidazole 30 mg/kg/d. Mice younger than 3wks were weighed daily and treated by oral gavage with antibiotic dissolved in PBS at concentrations necessary to deliver a volume of 10μl/g. After weaning, mice were administered antibiotics in drinking water supplemented with 9.6mg/L Equal sweetener (∼3.2mg/kg/d). All control mice used for survival experiments were treated with vehicle controls.

### Microbiome reconstitution

To rapidly reconstitute microbiota following antibiotic treatment for evaluation of the MAIT cell developmental window, each group received FMT gavage from age-matched donor mice. Cecum and colon microbiota were collected anaerobically as with preparation for the *in vitro* assay and 50μl/mouse of the filtered PBSc slurry was used for FMT. Mice treated in all other experiments were co-housed with age-matched mice for 1wk following antibiotic treatment for microbiome reconstitution.

### Mouse tissue processing

For the isolation of skin cells, ears were excised, split into ventral and dorsal halves, and placed dermal side down for 1h 45m incubation at 37°C in 500μL RPMI 1640 with 20 mM HEPES (Corning), 50 mM β-mercaptoethanol (Gibco), 1 mM sodium pyruvate (Corning) , 2 mM glutaGRO (Corning), 1 mM nonessential amino acids(Corning), 100 U/mL penicillin (Corning), and 100 mg/mL streptomycin (Corning) (Supplemented RPMI), 0.5mg/ml DNase I (Sigma-Aldrich) ,and 0.25 mg/ml of Liberase TL (Roche) (Digest Media). After incubation, 500μL Supplemented RPMI with 0.5mg/ml DNase I, 3% Fetal Bovine Serum (FBS) and EDTA was added to stop digestion. Digested tissue was then passed through a 70μm filter in Supplemented RPMI with 0.5mg/ml DNase I and 3% FBS (3% DNase), centrifuged, and resuspended in 500μL Supplemented RPMI with 10% FBS (10% RPMI).

For the isolation of lung cells, lungs were excised, diced, and incubated in Digest Media for 45m in a 37°C water bath with vortexing every 15 minutes. The digested lungs were passed through a 70-μm filter with 3% DNase, centrifuged, resuspended in 5 mL of 37% Percoll (Sigma-Aldrich), centrifuged, resuspended in 2mL RBC Lysis Buffer (BioLegend, 420302), incubated for 3min at room temperature, diluted with 3mL 3% DNase, centrifuged, and resuspended in 10% RPMI.

### Flow cytometry

Fluorophore-conjugated antibodies were purchased from BD Biosciences, BioLegend, or Invitrogen. 5-(2-oxopropylideneamino)-6-D-ribitylaminouracil (5-OP-RU)-loaded mMR1 and PBS57-loaded mCD1d tetramers were acquired from the NIH Tetramer Core Facility.^39^ For extracellular staining, cells were incubated in complete RPMI at room temperature for 1h. Cells were fixed for 1h and then stained intracellularly for transcription factors for 1h at 4°C using eBiosciences FOXP3/Transcription Factor Staining Buffer Set (Invitrogen, 00-5523-00). All staining mixes included TruStain FcX rat anti-mouse CD16/32 (BioLegend, 101320). Dead cells were removed by staining with LIVE/DEAD Fixable Blue Dead Cell Stain Kit (Invitrogen Life Technologies). Cytometric analysis was performed using a Cytek Aurora Spectral Flow Cytometer (Aurora Biosciences) with SpectroFlo software (Cytek). Data were processed using FlowJo (BD). MAIT cells were gated mMR1 tetramer^+^ TCRβ^+^ TCRγδ^−^ CD90.2^+^ CD45^+^ LIVE/DEAD Blue^-^; iNKT cells were gated as mCD1d tetramer^+^ mMR1 tetramer^−^ TCRβ^+^ TCRγδ^−^ CD90.2^+^ CD45^+^ LIVE/DEAD Blue^-^; CD4T cells were gated as FoxP3^-^ CD4^+^ CD8β^−^ mCD1d tetramer^−^ mMR1 tetramer^−^ TCRβ^+^ TCRγδ^−^ CD90.2^+^ CD45^+^ LIVE/DEAD Blue^-^; regulatory T cells (Treg) were gated as FoxP3^+^ CD4^+^ CD8β^−^ mCD1d tetramer^−^ mMR1 tetramer^−^ TCRβ^+^ TCRγδ^−^ CD90.2^+^ CD45^+^ LIVE/DEAD Blue^-^; CD8^+^ T cells were gated as CD8β^+^ CD4^−^ mCD1d tetramer^−^ mMR1 tetramer^−^ TCRβ^+^ TCRγδ^−^ CD90.2^+^ CD45^+^ LIVE/DEAD Blue^-^; and γδ T cells as TCRγδ^+^ TCRβ^−^ CD90.2^+^ CD45^+^ LIVE/DEAD Blue^-^.

### Francisella tularensis

*Francisella tularensis* subsp. *holarctica,* CDC Live Vaccine Strain (BEI Resources, NR-646) was streaked on cysteine heart agar. Single colonies were selected for growth in tryptic soy broth with 0.1g/L cysteine at 37°C. This was diluted to OD600 0.01 and grown to generate a CFU-OD curve. For *in vivo* experiments, the same inoculation and growth protocol was used, and the culture was grown until OD reached 0.6-0.8. CFU/ml was calculated and used for infection inoculates, which were diluted in PBS and confirmed by plating of inoculate prior to infection. Age- and sex-matched mice were weighed, anaesthetized with isoflurane, suspended, and inoculated retropharyngeally via reflexive aspiration. Weight was assessed daily. Mice were considered moribund upon weight loss of 25% or greater.

### Bacteroides thetaiotaomicron

*Bacteroides thetaiotaomicron,* VPI-5482 (ATCC, 29148) was grown anaerobically in BHI media supplemented with hemin and vitamin K overnight to stationary phase and plated to determine stationary CFU/mL. Broth culture was concentrated by centrifugation and resuspended in PBS for daily administration via oral gavage of 1 x 10^8^ CFU during antibiotic treatment.

### Enterococcus faecium

*Enterococcus faecium* (ATCC, BAA-2320) was grown anaerobically in BHI media supplemented with hemin and vitamin K overnight to stationary phase and plated to determine stationary CFU/mL. Broth culture was concentrated by centrifugation and resuspended in PBS for daily administration via oral gavage of 1 x 10^8^ CFU during antibiotic treatment.

### Statistical analysis

Statistical analyses were performed using Prism 10 software (GraphPad). Pairwise comparisons with 2 groups were performed using two-tailed unpaired Student’s *t* test; analyses with more than 2 groups were performed using ordinary one-way ANOVA with Šidák’s multiple comparison correction; *=*P* < 0.05, **=*P* < 0.01, ***=*P* < 0.001, and ****=*P* < 0.0001. “ns” = not significant. Survival statistics were performed using Mantel-Cox test.

## Supplemental information

Document S1. Figures S1-2

