## Supplemental figures S1-2 for "Antibiotic use in early life subsequently impairs MAIT cell-mediated immunity"

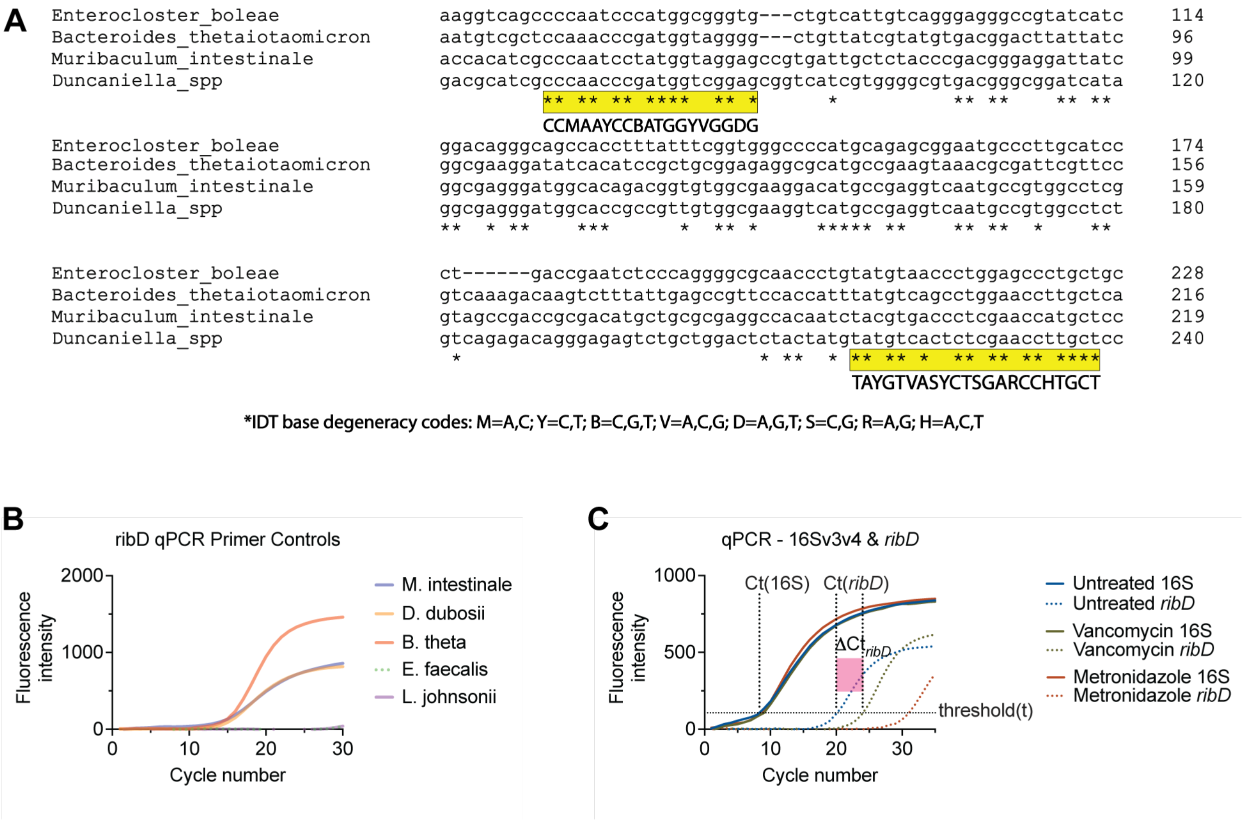


**Figure S1. Design of degenerate *ribD* primers**

(A) Multiple alignment and degenerate primer design for homology to *ribD* gene from diverse species isolated in our facility. (B) Evaluation of amplification of diverse *ribD* genes and for off-target binding by degenerate *ribD* primers. (C) Example data of qPCR of microbial DNA isolated following *in vitro* culture assay of weaning-age cecum and colon microbiota with or without antibiotics.

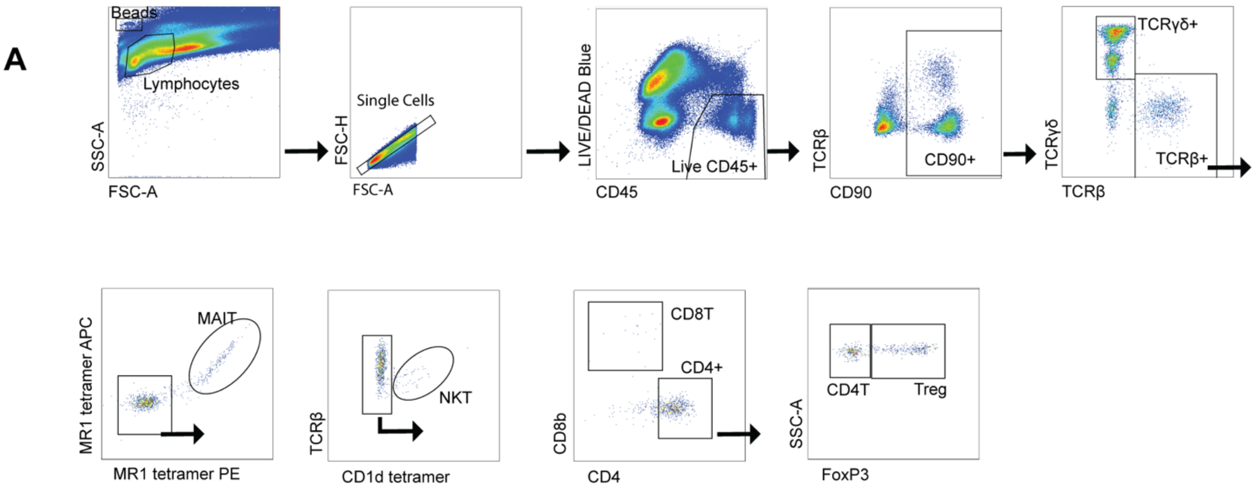


**Figure S2. Identification of lymphocytes by flow cytometry**

(A) Gating of lymphocytes using FlowJo. MAIT cells were gated mMR1 tetramer$^+$ TCRβ$^+$ TCRγδ$^-$ CD90.2$^+$ CD45$^+$ LIVE/DEAD Blue$^-$; iNKT cells were gated as mCD1d tetramer$^+$ mMR1 tetramer$^-$ TCRβ$^+$ TCRγδ$^-$ CD90.2$^+$ CD45$^+$ LIVE/DEAD Blue$^-$; CD4$^+$ T cells were gated as FoxP3$^-$ CD4$^+$ CD8β$^-$ mCD1d tetramer$^-$ mMR1 tetramer$^-$ TCRβ$^+$ TCRγδ$^-$ CD90.2$^+$ CD45$^+$ LIVE/DEAD Blue$^-$; regulatory T cells (Treg) were gated as FoxP3$^+$ CD4$^+$ CD8β$^-$ mCD1d tetramer$^-$ mMR1 tetramer$^-$ TCRβ$^+$ TCRγδ$^-$ CD90.2$^+$ CD45$^+$ LIVE/DEAD Blue$^-$; CD8$^+$ T cells were gated as CD8β$^+$ CD4$^-$ mCD1d tetramer$^-$ mMR1 tetramer$^-$ TCRβ$^+$ TCRγδ$^-$ CD90.2$^+$ CD45$^+$ LIVE/DEAD Blue$^-$; and γδ T cells as TCRγδ$^+$ TCRβ$^-$ CD90.2$^+$ CD45$^+$ LIVE/DEAD Blue$^-$.
